# Glutamatergic regulation of miRNA-containing exosome precursor trafficking and secretion from cortical neurons

**DOI:** 10.1101/2024.09.15.613153

**Authors:** Marcela Bertolio, Qiyi Li, Francesca E Mowry, Kathryn E Reynolds, Haichao Wei, Kyoeun Keum, Rachel Jarvis, Jiaqian Wu, Yongjie Yang

## Abstract

Neuronal exosomes are emerging secreted signals that play important roles in the CNS. Currently little is known about how glutamatergic signaling affects the subcellular localization of exosome precursor intraluminal vesicles (ILVs), microRNA (miR) packaging into ILVs, and *in vivo* neuronal exosome spreading. By selectively labeling ILVs and exosomes with GFP-tagged human CD63 (hCD63-GFP) in cortical neurons, we found that glutamate stimulation significantly redistributes subcellular localization of hCD63-GFP^+^ ILVs especially decreases its co-localization with multi-vesicular body (MVB) marker Rab7 while substantially promoting exosome secretion. Interestingly, glutamate stimulation only modestly alters exosomal miR profiles based on small RNA sequencing. Subsequent *in vivo* cortical neuronal DREADD activation leads to significantly more widespread hCD63-GFP^+^ area in hCD63-GFP^f/+^ mice, consistently supporting the stimulatory effect of glutamatergic activation on neuronal exosome secretion and spreading. Moreover, *in situ* localization of hCD63-GFP^+^ ILVs and secreted exosomes from specialized Hb9^+^ and DAT^+^ neurons were also illustrated in the CNS.

## Introduction

Extracellular vesicles (EVs, size 100-1000nm), primarily composed of plasma membrane-shed microvesicles and endosome-derived exosomes, are emerging to mediate important communication among heterogeneous CNS cell types (Blanchette and Rodal, 2020; Budnik et al., 2016; Wan et al., 2022). Neuron-secreted exosomes facilitate miRNA (miR) transfer into astrocytes or endothelial cells to subsequently up-regulate astroglial glutamate transporter GLT1 expression or to regulate brain vascular integrity (Men et al., 2019; Morel et al., 2013; Xu et al., 2017). Brain-derived neurotrophic factor (BDNF) induces miR sorting into neuron-secreted EVs which in turn increase excitatory synapse formation in recipient hippocampal neurons (Antoniou et al., 2023). Neuronal proteins Arc and PEG10 also share homology with retrotransposon Gag that can form EVs to facilitate transfer of mRNA between neurons (Pastuzyn et al., 2018; Segel et al., 2021). *In vivo* transmission electron microscopy, tracing, and proteomic experiments further demonstrated the involvement of exosomes in mediating transfer of transneuronally transported proteins (TNTPs) from retinogeniculate inputs to excitatory lateral geniculate nucleus (LGN) neurons and further to neurons in visual cortex (Schiapparelli et al., 2022). EV signaling was also found to mediate the transfer of Wingless (Wg) from synaptic boutons to the specialized muscle region sub synaptic reticulum (SSR) in the neuromuscular junction (NMJ) of fly larva (Koles et al., 2012) and prevents accumulation of ciliary cargo in C. elegans sensory neurons (Razzauti and Laurent, 2021). Additionally, neuronal exosomes have been shown to associate with various disease-causing mutant proteins in both neurodegenerative disease models and human CSF and plasma fluid (Delpech et al., 2019; Kim et al., 2022; You et al., 2023).

The understanding of the cellular regulation of exosome secretion from neurons by glutamatergic neurotransmission, the predominant signaling in the CNS, is at very beginning. Early *in vitro* studies showed that cultured cortical neurons secrete elevated exosomes upon KCL-induced depolarization (Faure et al., 2006) and GABAnergic blockade (Lachenal et al., 2011) based on increased detection of exosome protein markers. A number of other factors, ranging from disease-relevant proteins such as TDP43 and Tau to BDNF (Antoniou et al., 2023; Iguchi et al., 2016; Wang et al., 2017), also stimulate exosome secretion from cultured neurons. However, the ultracentrifugation isolation and exosome marker immunoblot-based quantification of secreted exosomes used in early studies tend to induce exosome damage and aggregation (Kim et al., 2022). These approaches are also less quantitative. In addition, although exosomes with putative neuronal surface markers are commonly found in human CSF in control and disease subjects (Hill, 2019), whether neuronal activity regulates *in vivo* neuronal exosome secretion and spreading remains unexplored. Similarly, whether the packaging of miRs, a major type of cargo in exosomes, into exosome precursor ILVs is influenced by neuronal glutamatergic activity is unknown, nor are whether the compartmental and subcellular distribution of ILVs. In the current study, by selectively inducing GFP-tagged ILV and exosome marker human CD63 (hCD63-GFP) in cortical neurons, we began to address these questions. We also illustrated *in situ* localization of ILVs and secreted exosomes from specialized dopaminergic and motor neurons in midbrain and spinal cord respectively, and how DREADD-mediated neuronal activation affects *in vivo* neuronal exosome spreading in the cortex.

## Materials and methods

### Animals

Human (h) CD63-GFP^f/f^ knock-in mice were previously generated and characterized by the lab (Men et al., 2019). Wild type (C57BL/6J background, #00664), DAT-Cre congenic (#006660), Hb9-Cre congenic (#006600) and Ai14-tdT^f/f^ congenic mice (#007914) were obtained from The Jackson Laboratory. Male and female mice were used in all experiments, and they were maintained on a 12-hour light/dark cycle with ad libitum access to food and water. All animal care and treatment procedures were strictly carried out in accordance with the NIH Handbook for the Care and Use of Laboratory Animals and the Guidelines for the Use of Animals in Neuroscience Research. The animal techniques utilized in this study were approved by the Tufts University’s Institutional Animal Care and Use Committee (IACUC).

### Drug administration

Clozapine-N-oxide (CNO) was reconstituted at a concentration of 10 mg/mL in DMSO and subsequently diluted in saline to a final concentration of 0.5 mg/mL as a working dosage. The mice received CNO via intraperitoneal injection (i.p.) at a dose of 0.3 mg/kg, two weeks following stereotaxic surgery. After injection, the mice were kept alive for 1 hour to assess neuronal activation, or 3 hours for ILVs/exosome spreading analysis.

### Primary neuronal culture and transfection

For neuronal primary cultures, cortical neurons were isolated from embryonic day 14–16 mouse brains. Cerebellum, olfactory bulbs, meninges, and hippocampus were removed from each brain. Cortices were placed into 0.05% trypsin solution for 10 min in a 37°C bead bath. The enzymatic reaction was stopped by addition of neuron plating medium (NPM), composed of neurobasal medium, 2% B27 neurobasal supplement, 2 mM glutamine, 1% of 100x GlutaMAX, 1% penicillin– streptomycin and 5% fetal bovine serum. The tissue was gently dissociated by pipetting with a 1 mL micropipette and dissociated cells were filtered through a 70 μm strainer to collect the neuron cell suspension. Freshly prepared neurons were then plated on 10 cm cell culture on precoated 12 mm diameter coverslips (Neuvitro Corporation) (2,5 x 10^4^ cells) placed in 24-well plates. The cultures were kept in NPM for 24 hr and then the media was replaced by neuron growth media (NGM) composed of neurobasal medium, 2% B27 neurobasal supplement, 2 mM glutamine, 1% of 100x GlutaMAX, and 1% penicillin–streptomycin. Neuronal cultures have typically less than 2% of glial cells based on GFAP immunostaining. To induce the expression of hCD63-GFP, AAV9-CaMKII-0.4.Cre (final concentration 1 x 10^10^ genome copy (gc)/mL; Addgene, #105558) was added to primary neuronal cultures at 1 day *in vitro* (DIV). Primary neurons transfection on precoated coverslips with Cy5-miR-124-3p (final conc. 25 nM) was performed with DharmaFECT Reagent (Thermo Fisher) following manufacturer’s instructions.

### Glutamate treatment

Prior to each experiment, a fresh 10 mM solution of L-glutamic acid (Sigma) in PBS was prepared. At DIV 7 (10 cm dishes) or DIV 9 (12 mm coverslips), the media was removed, and the cortical neuron cultures were rinsed with sterile PBS. Neurons were treated with 100 μM glutamate or with an equal volume of sterile PBS in NGM for 4 hs at 37 °C. The neuronal conditioned media (NCM) and cells were collected and stored at −80 °C for further analysis.

### Immunostaining

For immunocytochemistry, hCD63-GFP^f/+^ primary cortical neurons cultured on coated coverslips were washed once with PBS and fixed with 4% paraformaldehyde (PFA) for 15 min. Coverslips were rinsed three times in PBS followed by a solution of 0.2% Triton X-100 in PBS for 5 min. Then, the fixed permeabilized cells were incubated in blocking buffer (5% normal goat or donkey serum (NGS or NDS) and 0.1% Triton X-100 in PBS) for 1 hour at room temperature. Primary antibodies for Rab7 (1:100, Cell Signaling #9367 or #95746), EEA1 (1:100, Cell Signaling #3288), Golga5 (1:100, Cell Signaling #A15768), mouse CD63 (1:50, Biolegend #143901) and βIII-tubulin (1:1000, R&D Systems #MAB1195) were incubated overnight at 4 °C in blocking buffer. After washing coverslips three times in PBS, corresponding secondary antibody in PBS (1:1000) was added for 2 hr at room temperature. The cells were rinsed three times in PBS before mounting with ProLong Gold antifade reagent with DAPI (Invitrogen).

For immunohistochemistry, the animals were subjected to deep anesthesia through intraperitoneal injection (i.p.) of a cocktail of Ketamine (100 mg/kg) and Xylazine (10 mg/kg) in saline, after which intracardial perfusion with 4% PFA in PBS was performed. The brains and spinal cords were dissected and kept in 4% PFA overnight at 4 °C, then cryoprotected by immersion in 30% sucrose for 48 hr. Tissues were embedded and frozen in Tissue-Tek® OCT Compound (Sakura, Tokyo Japan). Coronal sections (20 or 40 µm) were prepared with a cryostat (Leica HM525) and mounted on glass SuperFrost Plus slides (Fisher Scientific). For neuronal activity evaluation, slides were rinsed three times in PBS, then treated with blocking buffer (1% BSA, 5% goat-serum, and 0.2% Triton-X-100 in PBS) for 45 min at room temperature and incubated with primary antibody for c-Fos (1:100, Cell Signaling #2250) overnight at 4 °C in blocking buffer. After washing slides three times in PBS, corresponding secondary antibody was added for 2 hours at room temperature. The sections were rinsed three times in PBS before mounting with ProLong Gold antifade reagent with DAPI (#P36931, Invitrogen). For visualization of hCD63-GFP fluorescence, slides were washed three times in PBS before mounting with ProLong Gold antifade reagent with DAPI (#P36931, Invitrogen).

### Image acquisition and analysis

Imaging of hCD63-GFP neurons was performed using the Leica Falcon SP8 confocal laser scanning microscope (5-8 μm Z-stack, 0.5 μm step) magnified with ×63 oil-immersion objective. Images were imported into FIJI (ImageJ2 version 2.14.0/1.54f) for further analysis. For hCD63-GFP^+^ puncta distance quantification from nucleus, single neurons were isolated by drawing regions of interest (ROIs) of βIII-tubulin. Blue (DAPI) and green (GFP) channels were converted into binary images and a threshold value was applied for each one. To determine the X/Y coordinates of nucleus and hCD63-GFP^+^ particles, the center of mass was measured by selecting the Analyze Particle option. Using the Euclidean distance formula, the distance from hCD63-GFP^+^ particles to the nucleus center of mass was calculated. Distances were averaged per cell. Subcellular distribution of hCD63-GFP^+^ puncta was calculated as a percentage of the total thresholded GFP signal area found on an individual neuron, defined by the βIII-tubulin ROI. ROIs for soma and neurites were manually drawn and the GFP^+^ signal area was measured on each ROI. GFP co-localization analysis with EEA1, Rab7 or Golga5 was performed on every stack of single cells using the BIOP JACoP plugin. For this, ROI of βIII-tubulin was defined, and the channels of interest were thresholded. Then, the thresholded M1 Mander’s coefficients were obtained. The co-localization between Rab7 and Golga5, or Cy5-miR-124-3p with hCD63-GFP was calculated as described, except that the neuron ROI was based on the DIC channel.

For representative whole brain images, 10x objective magnification images were captured using a Keyence BZ-X microscope and stitched together, while c-Fos images were taken with the Zeiss Axio Imager at 10x magnification. Analysis was performed with FIJI. For the quantification of hCD63-GFP^+^ brain tissue area, GFP intensity was first thresholded and ROIs containing signal were manually drawn on each side of the brain sections and their areas were measured. c-Fos immunostaining was quantified by setting an intensity threshold and the Integrated Density was measured. The values obtained for Control and hM3Dq brain sides were then normalized with the corresponding averaged Control value, and the percentage change between hM3Dq and Control side was calculated for each animal.

DAT-Cre^+^hCD63-GFP^f/+^Ai14-tdT^f/+^ whole brain representative images at P40 were taken with the Keyence BZ-X microscope at 10x magnification and stitched together, while Zeiss Axio Imager at 10x magnification was used to image Hb9-Cre^+^hCD63-GFP^f/+^Ai14-tdT^f/+^ whole spinal cord representative image at E12. Confocal 40x magnification images were captured with the Nikon A1R (15-20 Z-stack, 0.5 μm step size) and were utilized for the co-localization analysis between hCD63-GFP and tdT.

### ImmunoEM microscopy

Mice were deeply anesthetized and transcardially perfused with ice-cold heparinized PBS followed by fixation with 4% PFA + 0.25% glutaraldehyde in PBS. Brains were dissected, post-fixed overnight at 4 °C, and transferred to PBS. 100 μm coronal sections were cut on a vibratome and collected in PBS. Human CD63-GFP^f/+^ cortical neurons cultured on precoated coverslips were fixed with 4% PFA + 0.25% glutaraldehyde in PBS. Brain sections and neuron samples were processed for 5 nm immunogold labeling of GFP (anti-turbo GFP, Evrogen #AB513) at the Harvard Medical School Electron Microscopy Core facility. Images were acquired on a JEOL 1200EX transmission electron microscope. The number of axon terminals and dendritic spines containing immunogold particles (threshold for positive labeling: >1 immunogold particle) vs. lacking immunogold particles within synapse-containing images were expressed as percentages.

### Exosome purification and nanoparticle tracking analysis

Exosomes from NCM (5 x 10^6^ cells/10 cm dish) were isolated by size exclusion chromatography using the IZON qEVoriginal/35 nm columns following manufacturer’s instructions. Prior to the isolation process, the NCM was first spun at 300 x g for 10 min to remove cell debris, 4000 x g for 10 min at 4 °C, and then concentrated with Centricon Plus-70 filter (10K MWCO, Millipore) at 3500 x g at 4 °C until a volume of 500 μl or less was obtained. The concentrated sample was then transferred into the IZON column for separation. Fractions containing exosomes (#7 to #9) were combined and concentrated through Amicon Ultra-4 filter unit, and the final volume was adjusted to 100 μl for all samples. Extracellular vesicle size and concentration were analyzed by nanoparticle tracking analysis using the ZetaView instrument (Particle Metrix) and corresponding software ZetaView (version 8.05.11 SP1) at the Boston Children’s Hospital Cell Function and Imaging Core, Harvard Digestive Diseases Center (Boston, MA). Firstly, 100 nm “Standard beads” (Particle Metrix GmbH) were used to calibrate the machine. The following settings were used for all measurements: 11 positions, 2 cycles, minimal brightness of 18, minimal size of 10 nm and trace length of 10 s. Sample dilutions were adjusted using milliQ water to a final volume of 1 mL.

### Immunoblotting

Anti-mouse CD63 (1:500, Abcam #ab217345), anti-human CD63 (1:100, Santa Cruz #sc-5275), anti-CD81 (1:400, Santa Cruz #sc-166029), anti-Alix (1:100, Santa Cruz #sc-53540), anti-turbo GFP (1:1000, Evrogen #AB513), anti-EEA1 (1:1000, Cell signaling #3288), anti-Rab7 (1:1000, Cell Signaling #9367), anti-Arc (1:10000, Proteintech #66550), and anti-GAPDH (1:10000, Proteintech #60004-1-Ig) primary antibodies were used. Neuronal pellets and exosome fractions were homogenized with RIPA buffer (Thermo Scientific). Protein inhibitor cocktail (P8340, Sigma) was added in a 1/100 dilution to this lysis buffer prior to sample homogenization. Total protein amount of cell lysates was determined by DC protein assay (Bio-Rad). Forty micrograms of neuronal lysate or equal volumes of exosomal lysate were prepared with sample buffer 4X (Bio-Rad) and β-mercaptoethanol, heated at 95 °C for 5 min and loaded on stain-free 4–15% gradient sodium dodecyl sulfate polyacrylamide gel electrophoresis gels (Bio-Rad). Separated proteins were transferred onto a Polyvinylidene difluoride (PVDF) membrane using the Trans-Blot Turbo system (Bio-Rad). The membrane was blocked with 5% non-fat dry milk in TBST (Tris buffered saline with 0.1% Tween 20) then incubated with the appropriate primary antibody overnight at 4 °C. On the following day, the membrane was exposed to HRP-conjugated goat anti-rabbit or anti-mouse secondary antibody (1:10000) diluted in TBST. Bands were visualized using the ChemiDoc MP imaging system (Bio-Rad) by ECL chemiluminescent substrate (Clarity Max Western ECL Substrate, Bio-Rad). Exposure time was optimized for detecting different proteins.

### RNA isolation and miR qPCR

Total RNA was extracted from exosome fractions and neuronal cell pellets using TRIzol® reagent (Invitrogen) by following the manufacturer’s instructions. RNA concentration was measured using the NanoDrop® ND-1000 UV-Vis Spectrophotometer. Equal quantities (10 ng) of each sample were used to convert the miRs into cDNA using the TaqMan MicroRNA Reverse Transcription Kit (Applied Biosystems) with specific primers for each individual miR (included in each TaqMan MicroRNA Assay) and control U6 small nuclear (sn) RNA (Applied Biosystems; used for neuronal samples only). qPCR was performed in a StepOnePlus Real-Time PCR system (Applied Biosystems) with matching miR probes (TaqMan MicroRNA Assay) and TaqMan 2x Universal PCR Master Mix. Within each individual sample, the abundance of each miR was calculated relative to miR-9-5p, the most abundant neuronal miR. To determine the relative enrichment of individual miR in neuronal exosomes over its neuronal abundance, an Exosome miR Enrichment index was calculated by dividing the exosome fold change of a particular miR (relative to miR-9-5p) by its corresponding neuronal fold change (relative to miR-9-5p).

### Small RNA sequencing and analysis

Small RNA sequencing was performed at Tufts University Core Facility Genomics Core (Boston, MA). The quantity and quality of isolated exosomal RNA was determined using the Agilent 5200 Fragment Analyzer system. Library preparation was done with the QIAseq miRNA Library Kit (Qiagen), and the Illumina NovaSeq 6000 instrument was used for sequencing. The small RNA-seq data was analyzed at the University of Texas Health Science Center at Houston, McGovern Medical School (Houston, TX). Data preprocessing involved the removal of adapter, index, and low-quality reads using cutadapt (DOI:10.14806/ej.17.1.200). Subsequently, small RNA with sizes ranging from 17 to 40 nucleotides were selected for further analyses. Alignment of these small RNA was performed against mature miRNA sequences of mice sourced from miRbase using bowtie (DOI: 10.1186/gb-2009-10-3-r25). Samtools (DOI: 10.1093/bioinformatics/btp352) sorted the aligned reads to mature miRNA sequences and generated read counts for each miRNA. Normalization of miRNA raw reads was performed using Deseq2 (PMID: 25516281) with default parameter. Differential gene expression analysis was conducted using DESeq2 with identification of significant differentially expressed miRs based on criteria including an adjusted p-value less than 0.05 and a fold change greater than 2.

### Bioinformatic prediction of miR/mRNA binding

Predicted mRNA targets of top detected miRs in exosomes were determined using TargetScan Mouse 8.0. Targets with >1 conserved site were considered. The list of predicted targets was cross-referenced with datasets from Zhang et al. 2014 (neurons, microglia, oligodendrocytes, and endothelial cells) and Morel et al. 2017 (adult astrocytes) to determine cell-type specific mRNA transcripts, using thresholds of FPKM>10 and >4-fold enrichment relative to all other cell types.

### Stereotaxic delivery of AAV virus

A mixture of AAV8-hSyn-DIO-hM3Dq-mCherry (1.5 μL, 3.20 × 10^13^ gc/mL) (Addgene, #44361) and AAV9-hSyn-Cre-hGH (0.5 μL, 2.30 × 10^13^ gc/mL) (Addgene, #105555) was stereotaxically injected into the motor cortex of the right hemisphere of hCD63-GFP^f/+^ mice (Bregma 0, M/L = 1.0 mm, D/V = −1.0 mm). A mixture of AAV9-hSyn-Cre (0.5 μL, 1 × 10^13^ gc/mL) and PBS (1.5 μL, 1X) was stereotaxically injected into the motor cortex of the left hemisphere of hCD63-GFP^f/+^ mice (Bregma 0, M/L = −1.0 mm, D/V = −1.0 mm). AAV8-Syn-DIO-hCD63-GFP virus (0.6 μL, 1.31 x 10^14^ gc/mL) was injected bilaterally in the ventral tegmental area (VTA) of DAT-Cre mice (A/V = −3.1 mm from Bregma, M/L = ± 0.75 mm, D/V = −4.7 mm). AAV8-Syn-DIO-hCD63-GFP virus was packaged by the Boston Children’s Hospital Viral core (Boston, MA). Mice were anesthetized with isoflurane vapor (Covetrus). Post-operative care included injections of buprenorphine according to the IACUC requirement. Animals were perfused 14 days after injections.

### Statistical analysis

Data sets were analyzed using GraphPad Prism 10.1.1 software (La Jolla, CA, USA) and presented as mean ± SEM. Data sets were compared using two-tailed Student’s t-tests or one- and two-way ANOVA with post-hoc Šídák post-hoc testing unless otherwise indicated. Statistical significance was tested at a 95% (P < 0.05) confidence level and P values were shown in each graph.

## Results

### Glutamatergic signaling regulates compartmental and subcellular trafficking of tetraspanin CD63 in cortical neurons

We previously generated hCD63-GFP^f/f^ mice to selectively and efficiently label secreted exosomes and their intracellular precursor ILVs in a cell-type specific manner by tagging the C-terminus of exosome surface marker human (h) CD63 with the GFP reporter (Men et al., 2019). AAV-CaMKII-Cre induced hCD63-GFP expression has a peri-nuclear localization pattern in neuronal soma (Fig. 1Aii) which is highly similar and well overlapped (> 70%) with endogenous mouse (m) CD63 immunoreactivity (Fig. 1Ai, Fig. 1B). For acute glutamate treatment (up to 4 hrs.), glutamate (100 μM), comparable to evoked synaptically released glutamate concentration in previous studies (Herman and Jahr, 2007), was added to cortical neuronal cultures at days *in vitro* 7 (DIV7) when glutamate receptors are evidently expressed (Ha et al., 2009). Glutamate stimulation of neurons induced minimal neuronal cell death within 4 hrs., consistent with previous report (Ha et al., 2009). The overlap between endogenous mCD63 and hCD63-GFP is only slightly affected by acute glutamate treatment for up to 4 hours (Fig. 1B). Similarly, glutamate stimulation induced no expression changes of hCD63, mCD63, GFP, and other ILV marker CD81 (Fig. 1C and D). Interestingly, glutamate stimulation significantly increased the mean distance between all hCD63-GFP^+^ puncta to the center of nucleus (Fig. 1Ei-ii and F), suggesting an increasingly scattered distribution of hCD63-GFP signals from neuronal soma to the whole neuron. Consistently, glutamate stimulation significantly reduced the total hCD63-GFP^+^ puncta area in soma while increasing that in neurites (Fig. 1G). As there are no obvious expression changes of hCD63 and GFP induced by glutamate stimulation (Fig. 1C and D), it is unlikely that increased scattering of hCD63-GFP^+^ puncta by glutamate is due to hCD63-GFP expression changes.

**Figure 1.**
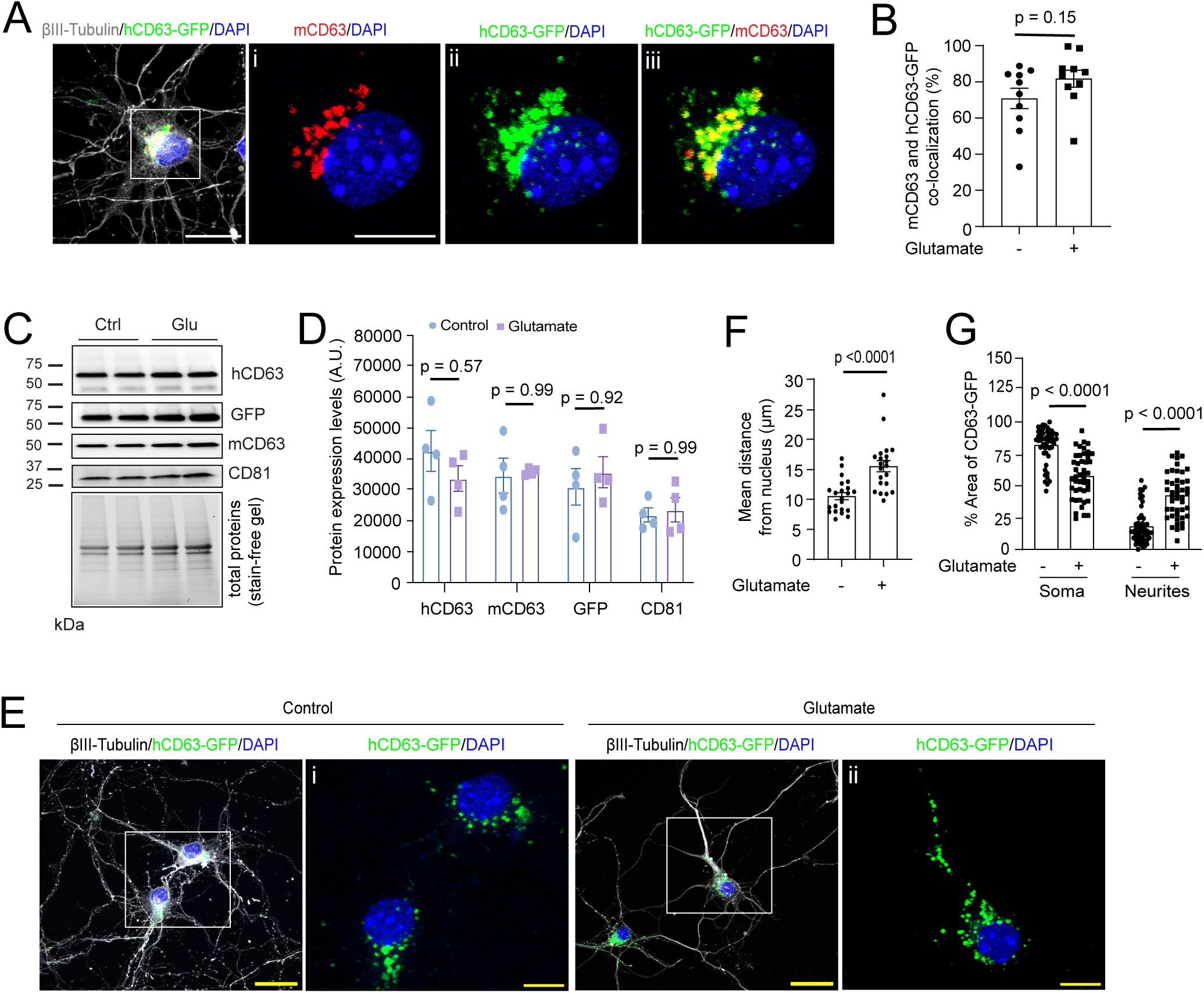
Glutamatergic signaling regulates compartmental localization of tetraspanin CD63 in cortical neurons. ***A.*** Representative confocal images of hCD63-GFP neurons with endogenous mouse CD63 immunoreactivity; Scale bar: 20 μm, 10 μm (i-iii); ***B.*** Quantitative analysis of co-localization comparison between control and glutamate treatment. n = 10-11 neurons/group; p values determined in two-tailed unpaired t test. Representative immunoblots (**C**) and quantification (***D***) of engineered exosome markers (hCD63, GFP) and endogenous exosome markers (mCD63, CD81) in whole neuronal cell lysates (WCL). n = 4 independent samples/group. Protein expression levels were normalized by total loaded proteins (stain-free gel). p values determined in one-way ANOVA with Tukey’s post hoc analysis. ***E.*** Representative confocal images of the subcellular distribution of hCD63-GFP^+^ puncta in cultured primary neurons following glutamate treatment. Scale bar: 30 μm and 10 μm (magnified view); ***F.*** Quantification of hCD63-GFP^+^ puncta distance from nucleus in control and glutamate treated neurons. n = 21-22 cells/group; p values determined from the two-tailed unpaired t test. ***G.*** Quantification of hCD63-GFP signal in soma and neurites, represented as a percentage of the total signal area found in the cell. n = 40-50 cells/group from 2 biological replicates; p values determined in the two-way ANOVA with post-hoc Šídák test.

As CD63 is a well-validated exosome surface marker (Kowal et al., 2016), its subcellular localization facilitates the examination of subcellular localization of ILVs, especially within the endosome/lysosome network (ELN). Currently, how CD63 is trafficked intracellularly in neurons remains unknown. Although it is a predicted plasma membrane protein like other tetraspanin family members (Pols and Klumperman, 2009), we observed no hCD63-GFP signal on the neuronal surface, nor with the mCD63 immunostaining (Fig. 1). To further characterize CD63 intracellular trafficking, we then performed immunostaining of EEA1 and Rab7, selective markers for early endosome and multivesicular body (MVB) respectively in hCD63-GFP induced primary neurons following glutamate stimulation (Fig. 2A). As the ER/Golgi network is able to directly deliver newly synthesized and modified proteins to endosomal compartments, we also performed immunostaining of Golga5 that selectively labels the Golgi network (Fig. 2A). Quantitative co-localization of hCD63-GFP with these organelle markers showed predominant co-localization of hCD63-GFP with MVB marker Rab7 (78%), but only ∼7% overlap with early endosome marker EEA1, ∼15% with Golgi marker Golga5, and <1% with DAPI (Fig. 2Ai-iii, i’-iii’, Fig. 2B), indicating that MVBs are the primary source of hCD63-GFP^+^ ILVs. Interestingly, glutamate stimulation had minimal effect on the co-localization of hCD63-GFP with EEA1, Golga5 (Fig. 2Ai’, iii’, Fig. 2B), and DAPI, but significantly reduced co-localization of hCD63-GFP with Rab7 (Fig. 2Aii’, Fig. 2B), even a significantly increased Rab7 protein expression upon glutamate stimulation was observed (Fig. 2C and D). Meanwhile, EEA1 protein expression was significantly decreased upon glutamate stimulation (Fig. 2C and D), which may correspond to the transition of early endosomes into MVBs by glutamate stimulation in the endosome pathway. As hCD63-GFP is localized on the surface of MVB-derived ILVs, its reduced co-localization with Rab7 by glutamate stimulation suggests that glutamate stimulation may induce release of ILVs from the MVB to extracellular space. This is in parallel with glutamate-induced increasingly scattered localization of hCD63-GFP in neurons that presumably suggests their trafficking near the plasma membrane for release.

**Figure 2.**
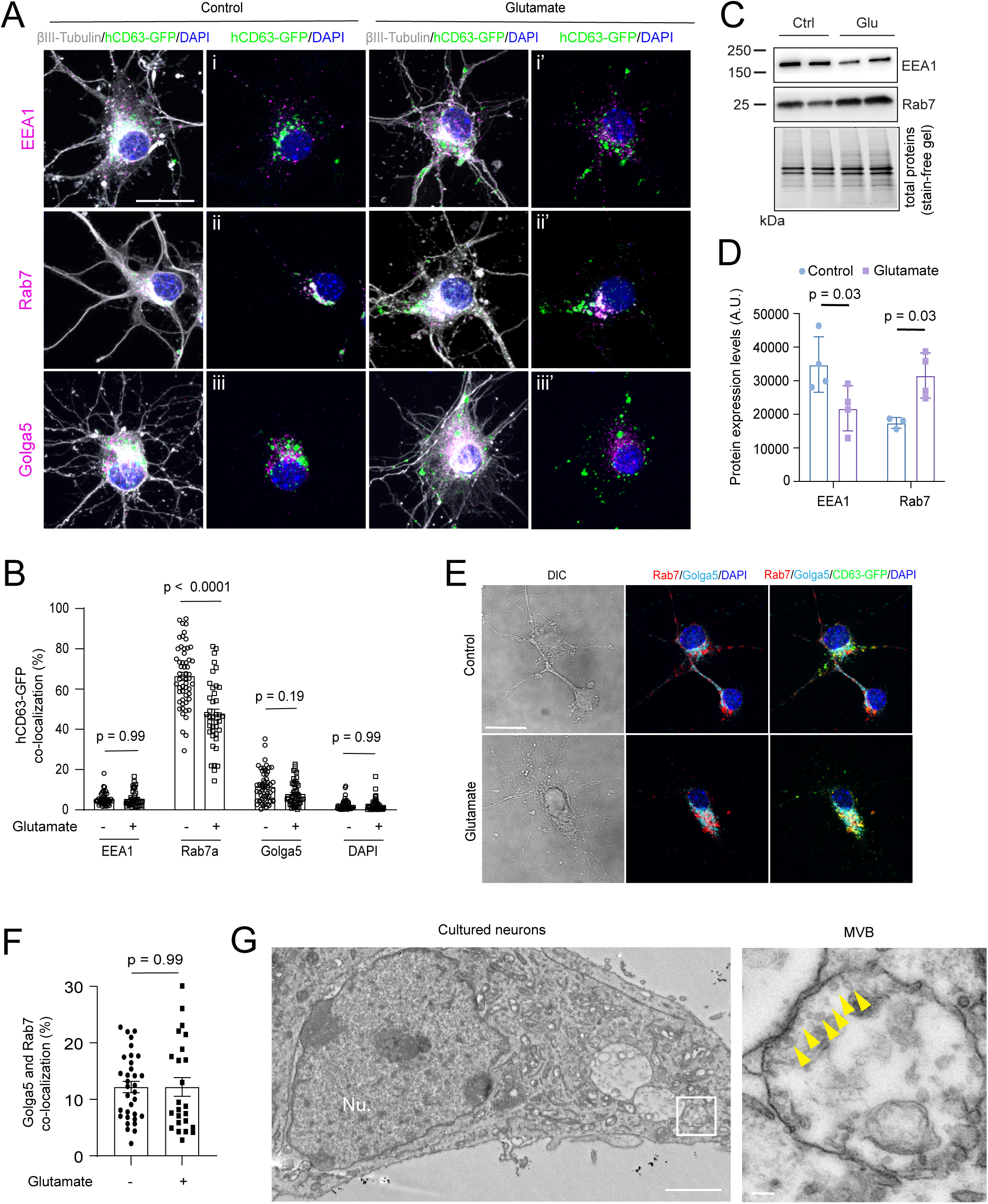
Glutamate stimulation affects the endosomal pathway and subcellular trafficking of hCD63-GFP^+^ vesicles. ***A.*** Representative confocal images of control and glutamate-treated (4hr) hCD63-GFP expressing neurons immunostained with EEA1 (early endosomes), Rab7 (late endosomes/MVBs), and Golga5 (Golgi apparatus) antibodies; Scale bar: 20 μm; ***B.*** Quantification of hCD63-GFP colocalization with individual organelle markers in control and glutamate treated neurons. n = 40-55 neurons/group from 2 biological replicates; p values determined from two-way ANOVA with post-hoc Šídák test; Representative immunoblots (***C***) and quantification (***D***) of endosomal markers EEA1 (early endosomes) and Rab7 (late endosomes/MVBs) in whole neuronal cell lysates (WCL). Protein expression levels were normalized by total loaded proteins (stain-free gel). n = 4 independent samples/group, two-way ANOVA with post-hoc Šídák test. ***E.*** Representative DIC and confocal images of Rab7 and Golga5 immunostaining in control and glutamate-stimulated hCD63-GFP^+^ neurons. Scale bar: 20 μm; ***F.*** Quantitative analysis of co-localization between Rab7 and Golga5 in control and glutamate-stimulated hCD63-GFP^+^ neurons. n = 24-33 neurons/group; two-tailed unpaired t test. ***G.*** Representative electron-microscopy image of a cultured cortical neuron and a magnified view of a MVB containing intraluminal vesicles (yellow arrows); Nu: nucleus; Scale bar: 100 nm.

In addition, significant higher co-localization of hCD63-GFP with Golga5 (∼15%, Fig. 2B) than co-localization of hCD63-GFP with EEA1 (∼7%, Fig. 2B) also suggests that a portion of newly synthesized hCD63-GFP traffics directly from Golgi networks to MVBs without being first targeted to the plasma membrane and subsequently endocytosed into early endosomes. Indeed, a similar percentage of co-localization between Rab7 and Golga5 (∼13%, Fig. 2E and F) was also observed. The co-localization between Golga5 and Rab7 (Fig. 2F) is not altered by glutamate stimulation. To overcome the resolution limit of light microscopy, we further performed GFP immunogold labeling and EM imaging on hCD63-GFP^+^ cultured neurons. Although GFP^+^ immunogold particles were observed (data not shown), the neuronal morphology was severely damaged during the immunoEM procedure to recognize the MVB structure. On the other hand, EM imaging without GFP immunogold labeling from cultured neurons found clear inwardly budding small vesicles, the ILVs, from the MVB (yellow arrows, enlarged view, Fig. 2G), clearly demonstrating MVB-derived exosome biogenesis.

### Glutamatergic signaling promotes exosome secretion from cortical neurons and modestly alters miRNA (miR) profile in neuronal exosomes

To directly examine how glutamate stimulation alters neuronal exosome secretion, we collected neuron-conditioned medium (NCM) 4h following glutamate treatment. Neuronal exosomes were isolated by size exclusion chromatography (SEC) and their size and quantity were determined by ZetaView nanoparticle tracking analysis (NTA). Exosomes secreted from control and glutamate-stimulated neurons were highly similar in size distribution (Fig. 3A). Interestingly, >98% vesicle size is typical of exosome size (50-200 nm) but not larger microvesicles, consistent with the predominant MVB localization of hCD63-GFP in neurons, suggesting that exosomes, but unlikely MVs, are the predominant secreted EVs from neurons. Glutamate stimulation increased exosome numbers in almost all size ranges (Fig. 3A), with a 45% increase in the total number of exosomes secreted from glutamate-stimulated neurons compared to control neurons (Fig. 3B). Subsequent immunoblotting detected GFP and hCD63, as well as several other well-validated exosome markers mCD63, CD81, and Alix (Fig. 3C), confirming that hCD63-GFP is indeed localized on the surface of exosomes, together with endogenous mCD63, to be secreted from neurons. Interestingly, although Arc has been shown to form a viral capsid-like structure to package its own mRNA(Pastuzyn et al., 2018), we detected no Arc in neuronal exosomes (Fig. 3C), possibly due to the heterogenous nature of secreted vesicles from neurons.

**Figure 3.**
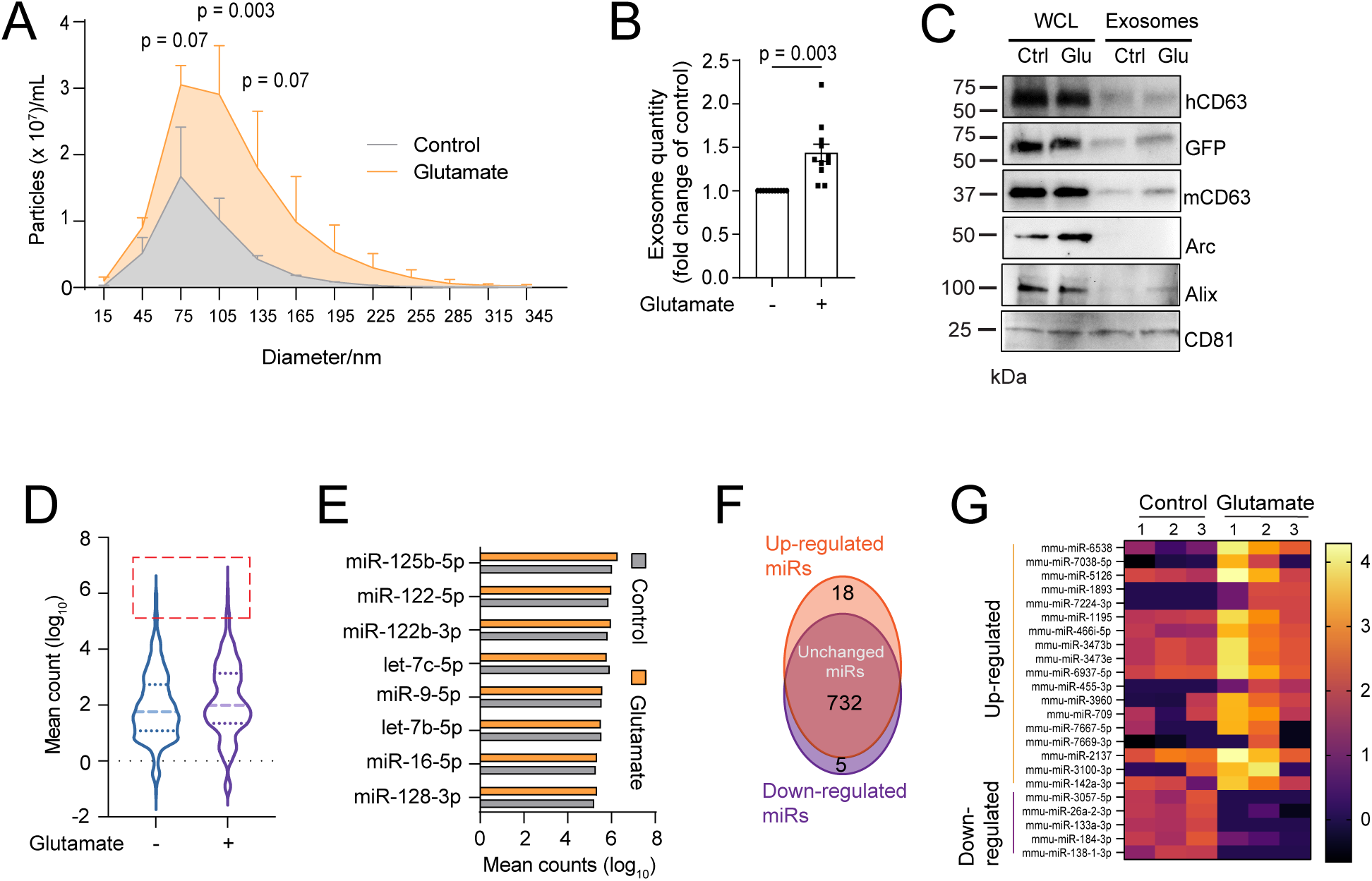
Glutamate stimulation significantly promotes exosome secretion from cortical neurons but modestly alters miRNA (miR) profile in neuronal exosomes. ***A.*** Representative size distribution of exosomes isolated from conditioned medium of control and glutamate treated cultured primary neurons. n = 3 replicates; p values determined from two-way ANOVA with post-hoc Šídák test. ***B.*** Relative fold change (over untreated control) of NTA measured total secreted exosomes from control and glutamate treated neurons. n = 11 individual replicates/group; p values determined from two-tailed unpaired t test. ***C.*** Representative immunoblots of engineered GFP tag, human CD63, and typical exosome protein markers from hCD63-GFP^+^ whole cell lysate (WCL) and purified exosomes. ***D.*** Violin plot depicting the distribution of all detected miRNAs in exosomes isolated from control and glutamate-treated neurons. The mean count from each group (3 replicates/group) was converted (log_10_). Highly expressed miRNAs in both conditions are indicated by the red rectangle and graphed in ***E***. ***F.*** Venn diagram of glutamate-induced up-, down-regulated, and unchanged exosomal miRs determined by small RNA-seq analysis; Up- and down-regulated miRs were identified based on padj < 0.05 and fold change > 2; ***G.*** Heatmap showing the up-regulated and down-regulated exosomal miRNAs (padj < 0.05, log_10_ mean counts) from cultured primary cortical neurons after glutamate exposure. n = 3 replicates/group.

Although miRs are considered a major category of neuronal exosomal cargoes, whether glutamatergic signaling alters miR content in neuronal exosomes remains unexplored. We therefore sequenced small RNAs isolated from exosomes purified from control and glutamate-treated neuronal cultures. The overall mean counts and number of detected miRs were highly similar in both groups (Fig. 3D). In addition, the very top detected miRs (mean counts > 10^5^), miR-125b-5p, miR-9-5p, miR-128-3p etc., were also highly conserved in both control and glutamate-stimulated neuronal exosomes with no differences in mean counts (Fig. 3E). Out of ∼ 750 total detected miRs, 18 and 5 were significantly (p < 0.05, FC > 2) up- and down-regulated in neuronal exosomes by the glutamate stimulation, respectively (Fig. 3F and G).

We further examined the abundance of top detected exosomal miRs and miR-124-3p (also within 10% detected miRs in exosomes) based on our previous studies (Men et al., 2019; Morel et al., 2013), in neurons and neuronal exosomes by quantitative PCR to determine their relative enrichment in exosomes. Although miR-9-5p was highly expressed and far more abundant than miR-124-3p in neurons (Fig. 4A), it was less proportionally abundant in exosomes (Fig. 4B) compared to miR-124-3p, indicating that these miRs were differentially enriched in exosomes regardless their neuronal abundance (Fig. 4C). These results support a selective sorting mechanism in packaging certain miRs into ILVs in neurons. Interestingly, glutamate stimulation had no major impact on miR abundance in neurons and exosomes (Fig. 4A and B) with similar miR enrichment in neuronal exosomes (Fig. 4C).

**Figure 4.**
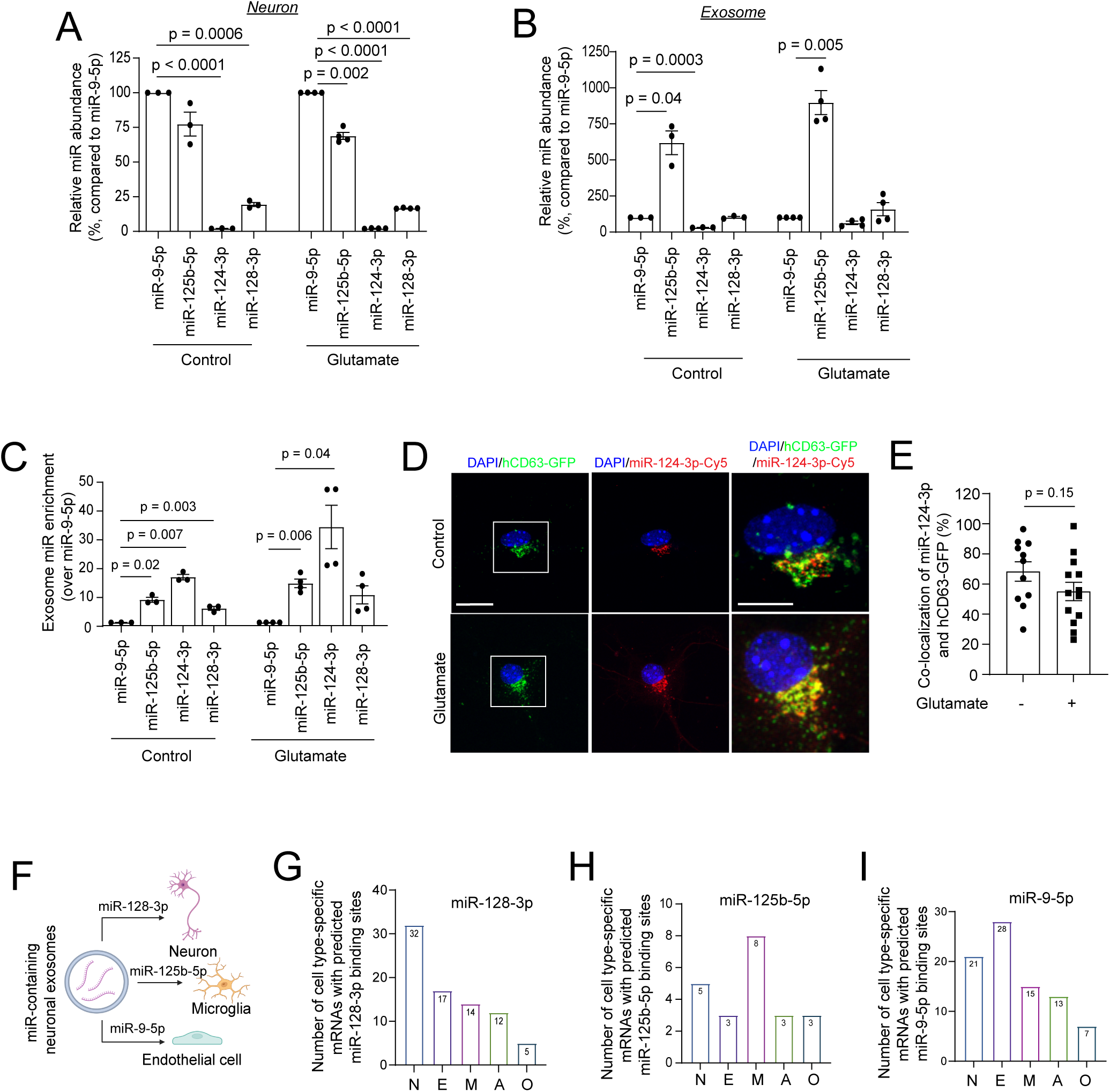
Neuronal miRs are differentially enriched in neuronal exosomes with minimal impact by glutamate stimulation. Relative miR abundance (%, in comparison to miR-9-5p) in neurons (***A***) and exosomes (***B***) in both control and glutamate stimulated neurons. n = 3-4 independent cultures/group; p values determined from repeated measures one-way ANOVA with Dunnett’s multiple comparisons test. ***C.*** Relative enrichment of miRs into exosomes (fold change in comparison to miR-9-5p) from control and glutamate stimulated neurons. miR abundance in neurons and exosomes was determined using qPCR and individually normalized to each sample’s miR-9-5p level which is most abundant in neurons. A higher enrichment index indicates increased packaging of miR into exosomes. n = 3-4 independent cultures/group; p values determined from repeated measures one-way ANOVA with Dunnett’s multiple comparisons test. ***D.*** Representative confocal images of transfected Cy5-miR-124-3p in control and glutamate-treated hCD63-GFP^+^ neurons; Scale bar: 20 μm and 10 μm (magnified view); ***E.*** Quantification of co-localization between Cy5-miR-124-3p and hCD63-GFP^+^ puncta in control and glutamate treated neurons. n = 11-13 neurons/group; p value determined from the two-tailed unpaired t test. Schematic diagram (***F***) of potential targeted delivery of miR-128-3p, miR-125b-5p, and miR-9-5p to neurons (***G***), microglia (***H***), and endothelial cells (***I***), respectively, based on the number of cell type-enriched predicted targets showed in the graphs. Predicted mRNA targets of top detected miRs in exosomes were determined using TargetScan Mouse 8.0. Targets with >1 conserved site were considered. Cell-type specific mRNA transcripts were determined with thresholds of FPKM>10 and >4-fold enrichment relative to all other cell types.

To examine whether miR cargoes are indeed packaged into ILVs inside neurons, we transfected hCD63-GFP^+^ primary cortical neurons with Cy5-labeled miR-124-3p, one of the highly detected miRs in neuronal exosomes and is functionally significant in mediating neuron to astroglia communication (Men et al., 2019; Morel et al., 2013), to examine its co-localization with hCD63-GFP^+^ ILVs inside neurons. Transfected Cy5-miR-124 was primarily localized within neuronal soma (70%) near the nucleus (Fig. 4D), similar to the subcellular distribution of hCD63-GFP. Indeed, a high percentage (68%) co-localization between Cy5-labeled miR-124-3p and hCD63-GFP^+^ ILVs was observed in neurons (Fig. 4E), confirming its substantial packaging into ILVs. Glutamate stimulation only slightly reduces co-localization of Cy5-miR-124-3p with hCD63-GFP (p = 0.15, Fig. 4E), consistent with the modest effect of glutamate on exosomal miR profile observed above. As exosomal miRs have been shown to mediate a number of functions intercellularly, including in the CNS, we then analyzed whether certain top detected miRs preferentially target on mRNAs that are selectively expressed or enriched in specific CNS cell types (Fig. 4F) based on previously published CNS cell type transcriptome databases (Morel et al., 2017; Zhang et al., 2014). Our analysis showed that miR-128-3p is predicted to bind to the largest number of neuron-selective mRNAs (Fig. 4G) and miR-125b-5p preferentially targets on microglial mRNAs (Fig. 4H), while miR-9-5p preferentially targets on endothelial cell mRNA (Fig. 4I). The selective targeting of cell-type specific mRNAs by different highly detected exosomal miRs suggests potentially pleiotropic functions of neuronal exosomes.

### *In vivo* neuronal excitation promotes exosome spreading from cortical neurons

Whether neuronal activity influences spreading of neuronal exosomes *in vivo* remains unexplored. We decided to address this question by employing the DREADD-based chemogenetic approach to precisely modulate activity of targeted neurons following administration of DREADD selective agonist CNO. We were particularly interested in the effect of neuronal excitation on exosome secretion, which can be more clearly assessed than inhibition from targeted neurons. The overall injection paradigm and experimental groups are illustrated in Fig. 5A, in which AAV-hSyn-Cre or a mixture of AAV-hSyn-Cre and AAV-DIO-hM3Dq-mCherry are separately and focally injected into the motor cortex of hCD63-GFP^f/+^ mice (Fig. 5A). In this paradigm, focal injections of equal AAV-hSyn-Cre virus leads to parallel hCD63-GFP induction from both cortical sides, while the mCherry-tagged hM3Dq receptor is only induced in neurons on one cortical side injected with the mixture of AAV-hSyn-Cre and AAV-DIO-hM3Dq-mCherry. As a result, hCD63-GFP^+^ areas, indication of ILVs and secreted exosomes from neurons, can be visualized and compared following DREADD-mediated neuronal activation.

**Figure 5.**
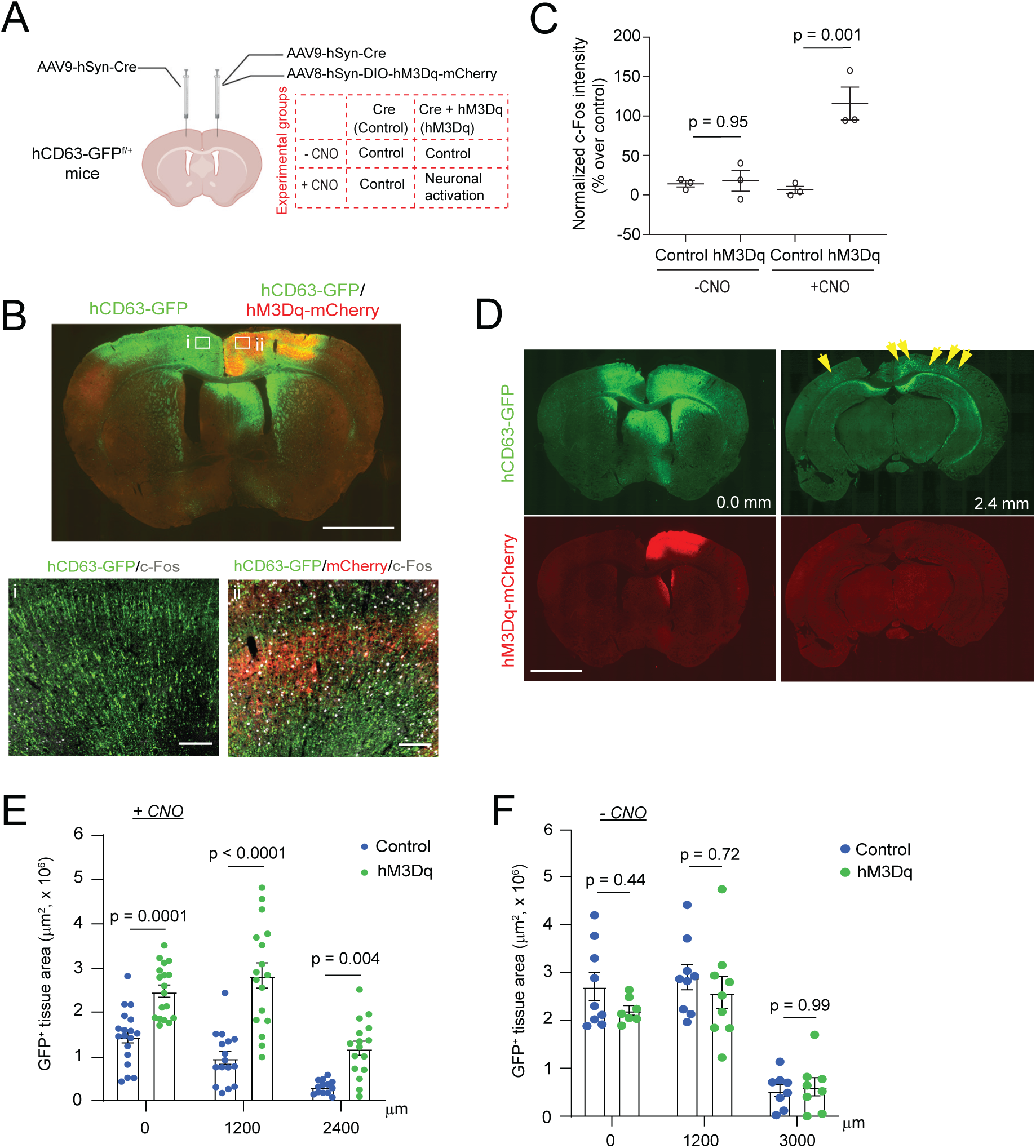
*In vivo* DREADD-mediated neuronal excitation promotes exosome spreading from cortical neurons. ***A.*** Experimental diagram and groups of bilateral and focal AAV injections into the motor cortex of hCD63-GFP^f/+^ mice. One cortical side received a mixture of AAV8-DIO-hM3Dq-mCherry and AAV9-hSyn-Cre virus, and the contralateral cortical side was injected with only AAV9-hSyn-Cre virus as a control. Experimental groups were also shown. ***B.*** A representative image of the bilateral injection and magnified view of both cortical sides (i: control side; ii: hM3Dq side, scale bar = 1 mm and 100 μm (magnified views) following CNO administration and c-Fos immunostaining. ***C.*** Quantification of c-Fos intensity from control and hM3Dq^+^ cortical sides of injected mice with or without CNO administration, represented by the percentage change of normalized c-Fos signal intensity relative to the control side. n = 3 mice, 3-5 fields/5 brain sections/mouse. P values determined from one-way ANOVA. ***D.*** Representative images (10x) from near the injection site (0.0 mm) showing the expression of hCD63-GFP and hM3Dq-mCherry and from the distant section (2.4 mm from the injection site) showing the continuous expression of hCD63-GFP (with no visible mCherry signals) following CNO administration; Scale bar: 1 mm; Quantification of hCD63-GFP^+^ brain area at different distance from the injection site (0.0, 1.2, and 2.4 mm) from CNO-administered (+CNO, (***E***) and CNO free (-CNO, (***F***) mice. Note the sections were 1.2 and 3 mm from the injection site in -CNO groups. n = 3 images/distance/mouse, 6 mice/group; p values determine from two-way ANOVA with post-hoc Šídák test.

The AAV-hSyn-Cre and AAV-DIO-hM3Dq-mCherry virus titer and CNO dose/duration have been tested to achieve optimized hCD63-GFP and hM3Dq-mCherry induction and neuronal excitation. As expected, hM3Dq-mCherry was found only on the side with the AAV-hSyn-Cre/AAV-DIO-hM3Dq-mCherry injection (Fig. 5Bii), while hCD63-GFP signals were observed on both cortical sides (Fig. 5Bi-ii). Subsequent c-Fos immunoreactivity also showed substantial and selective increase, an indication of widespread neuronal excitation, only on the AAV-hSyn-Cre/AAV-DIO-hM3Dq-mCherry-injected cortical side with CNO administration (Fig. 5Bii, Fig. 5C) but not on the AAV-hSyn-Cre-injected side or without CNO (Fig. 5C), confirming the specific excitation effect of hM3Dq/CNO on cortical neuronal activity. Subsequent quantification of hCD63-GFP^+^ area found a consistently and significantly greater hCD63-GFP^+^ tissue area at both proximal (0.0 mm, Fig. 5D) and distant (2.4 mm, Fig. 5D) sections from the injection site in the AAV-hSyn-Cre/AAV-DIO-hM3Dq-mCherry-injected cortical side compared to AAV-hSyn-Cre-injected cortical side following CNO administration (Fig. 5E), but not without CNO administration (Fig. 5F). These *in vivo* DREADD-mediated neuronal excitation results are consistent with glutamate-stimulated exosome secretion in cultured cortical neurons, further supporting the notion that increased neuronal excitation promotes exosome secretion and spreading.

### *In vivo* illustration of intracellular hCD63-GFP^+^ ILVs and extracellular exosomes from specialized neuronal subtypes and regions in the CNS

*In vivo* illustration of neuron-secreted exosomes in the CNS remains very limited. By generating cell-type specific and inducible ILV/exosome reporter mice, we previously began to demonstrate the *in situ* localization of exosomes secreted from forebrain (CaMKII^+^) excitatory neurons in adult mice with confocal and immunoEM imaging (Men et al., 2019). Given the presence of highly diverse neuronal subtypes in the mammalian CNS, we set out to explore *in situ* localization of ILVs/exosomes from other representative neuronal subtypes such as Homeobox transcription factor (Hb9^+^) motor neurons (MNs) in the spinal cord and dopamine transporter (DAT^+^) dopaminergic neurons from midbrain. These neuronal subtypes were chosen based on their predominant ventral and rostral localization in the CNS. We first generated Hb9-Cre^+^Ai14-tdT^f/+^hCD63-GFP^f/+^ and DAT-Cre^+^Ai14-tdT^f/+^hCD63-GFP^f/+^ triple positive mice to selectively label spinal cord motor neurons and midbrain dopaminergic neurons, respectively, so that hCD63-GFP^+^ alone signals, presumably extracellular exosomes, and hCD63-GFP^+^tdT^+^ signals, an indication of intracellular ILVs, can be better separated. Spinal cord and midbrain sections from different ages of Hb9-Cre^+^Ai14-tdT^f/+^hCD63-GFP^f/+^ and DAT-Cre^+^Ai14-tdT^f/+^hCD63-GFP^f/+^ mice were prepared, and representative images were collected respectively with confocal microscopy. Extracellularlly localized hCD63-GFP signals outside of tdT^+^ neuronal cell body, presumably secreted exosomes, were found as early as E12 in the spinal cord of Hb9-Cre^+^Ai14-tdT^f/+^hCD63-GFP^f/+^ mice (white arrows, Fig. 6A) and E18 in substantia nigra of DAT-Cre^+^Ai14-tdT^f/+^hCD63-GFP^f/+^ mice (white arrows, Fig. 6B). Quantification of hCD63-GFP^+^ alone and hCD63-GFP^+^tdT^+^ co-localized signals found consistent and substantial secreted exosomes, indicated by 18-20% hCD63-GFP^+^ alone signals, from Hb9^+^ and DAT^+^ neurons across early postnatal developmental stages (Fig. 6C).

**Figure 6.**
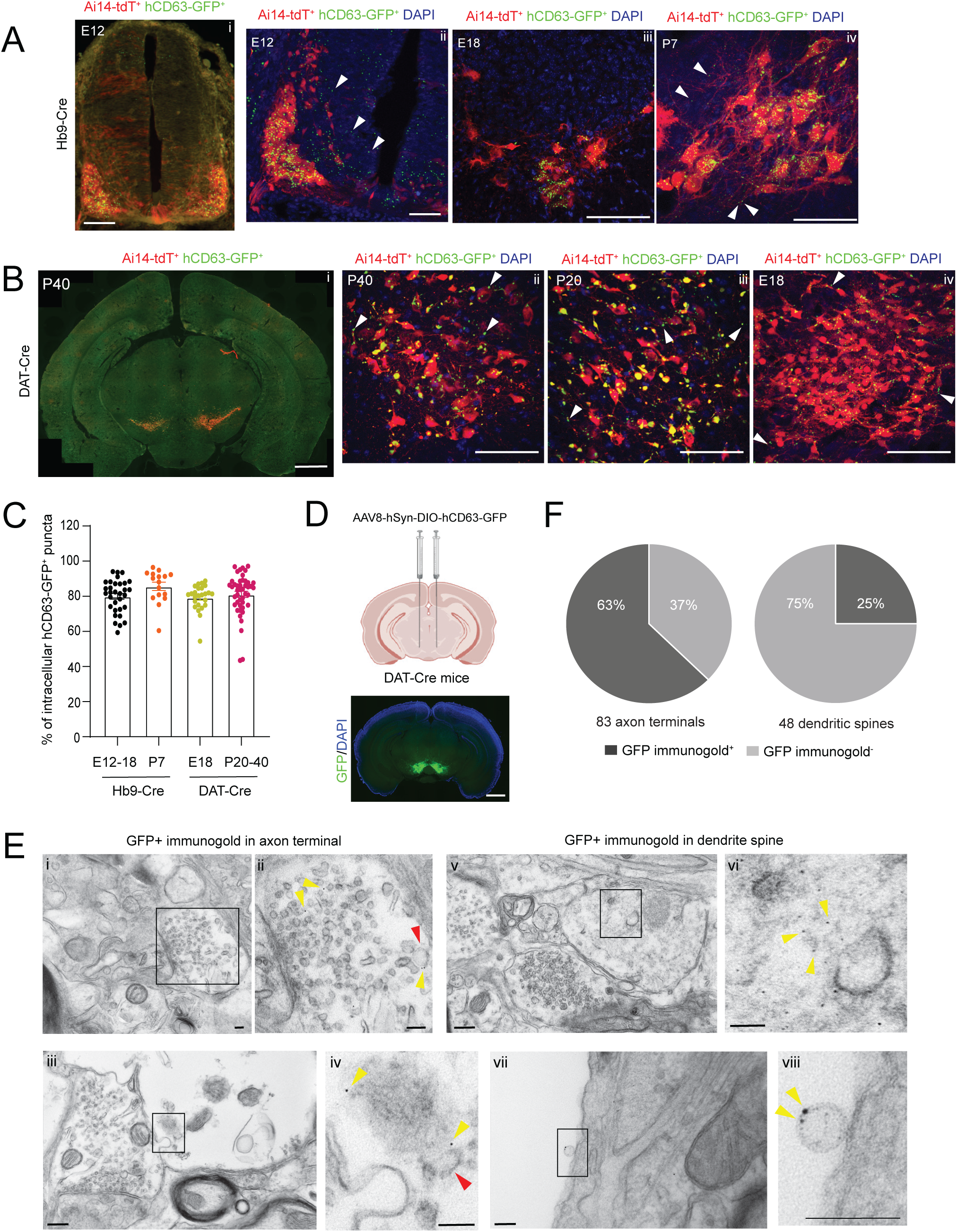
*In vivo* illustration of intracellular hCD63-GFP^+^ ILVs and extracellular exosomes from specialized neuronal subtypes and regions in the CNS. ***A.*** Representative confocal images of spinal cord sections of Hb9-Cre^+^hCD63-GFP^f/+^Ai14-tdT^f/+^ mice showing tdT^+^ motor neurons with intracellular (tdT^+^GFP^+^) and extracellular hCD63-GFP signals (tdT^−^GFP^+^, white arrows) at prenatal and early postnatal stages. Scale bar: 200 μm (i), 50 μm (ii), 100 μm (iii-iv); ***B.*** Representative confocal images of midbrain sections of DAT-Cre^+^hCD63-GFP^f/+^Ai14^f/+^ mice showing tdT^+^ dopaminergic neurons and intracellular (tdT^+^GFP^+^) and extracellular hCD63-GFP signals (tdT^−^GFP^+^, white arrows) at prenatal and early postnatal stages. Scale bar: 2 mm (i), 100 μm (ii-iv); ***C.*** Quantitative analysis of co-localization between hCD63-GFP and tdT at embryonic and early postnatal stages in spinal cord and midbrain sections from Hb9-Cre^+^hCD63-GFP^f/+^Ai14-tdT^f/+^ and DAT-Cre^+^hCD63-GFP^f/+^Ai14^f/+^ mice; n=18-31 sections, 3-5 images/4 sections/3-4 mice/group. ***D.*** The Diagram of bilateral injections of AAV9-hSyn-DIO-hCD63-GFP virus into the VTA of DAT-Cre^+^ mice. A representative image of selective induction of hCD63-GFP at the VTA from AAV9-hSyn-DIO-hCD63-GFP-injected DAT-Cre^+^ mice. Scale bar: 1 mm; ***E.*** Representative immunoEM images of hCD63-GFP^+^ intracellular signals and vesicles in axonal terminal (i-iv), dendrites (v-vi), blood vessel (vii-viii) from VTA of AAV9-hSyn-DIO-CD63-GFP-injected DAT-Cre mice. Scale bar: 100 nm. ***F.*** Pie charts of the percentage of axons or dendrites with GFP^+^ immunogold particles examined. n = 83 axon terminals and 48 dendrite spines respectively from 35 images of 2 mice.

To further illustrate hCD63-GFP labeling near synapses and blood vessels with increased resolution, we generated AAV-DIO-hSyn-hCD63-GFP virus and performed the stereotaxic injection into the ventral tegmental area (VTA) of adult DAT-Cre^+^ mice (Fig. 6D), which helps achieve increased hCD63-GFP signals to facilitate GFP immunogold labeling on midbrain sections. We first injected AAV-DIO-hCD63-GFP into the VTA and confirmed effective and focal induction of hCD63-GFP on DAT-Cre mice (Fig. 6D). As hCD63-GFP is only induced from DAT^+^ neurons in DAT-Cre mice, hCD63-GFP^+^ signals and labeled vesicles indicate intracellular ILVs and secreted exosomes from DAT^+^ neurons. In contrast to our previous observations from CaMKII^+^ excitatory neurons in forebrain (Men et al., 2019), hCD63-GFP^+^ immunogold signals were commonly (63% of all axon terminals examined, Fig. 6F) found in pre-synaptic axon terminals (Fig. 6Ei and enlarged view in ii, yellow arrows). Interestingly, the hCD63-GFP^+^ vesicle (red arrow, Fig. 6Eii) is substantially larger in size (100 nm) than typical synaptic vesicle size (20-30 nm) near the synaptic cleft. Excitingly, we also observed vesicular budding from the plasma membrane of the axon terminal (Fig. 6Eiii-iv) with a hCD63-GFP^+^ vesicle (50-100nm size, red arrow, Fig. 6Eiv) outside of the axon terminal near the budding site, supporting the secretion of hCD63-GFP^+^ vesicle from the axonal terminal. Dendritic labeling of hCD63-GFP was also observed (yellow arrows, Fig. 6Ev-vi, Fig. 6F). In addition, a hCD63-GFP^+^ vesicle with surface hCD63-GFP labeling (yellow arrows) was also found near the lumen of the blood vessel (Fig. 6Evii-viii). Together, these immunoEM results clearly support the widespread presence of secreted exosomes from DAT^+^ neurons in the VTA.

## Discussion

Our current study examined the cellular regulation of ILV biogenesis by glutamate stimulation in the endosome by following the subcellular trafficking of ILV and exosome surface marker CD63 in neurons. Our co-localization analysis between hCD63-GFP and early endosome and MVB markers, as well as EM imaging of cultured neurons, showed that ILVs are primarily localized in MVBs and are actively budded from MVBs. In addition, our quantitative measurement of exosome numbers also showed that glutamate stimulation significantly increases secreted exosome quantity, consistent with the significantly reduced MVB localization of hCD63-GFP^+^ ILVs by glutamate stimulation. It is noteworthy that our quantitative measurement of exosome numbers overcomes previous drawbacks of determining exosome quantity based on exosome protein marker levels, which may only indicate exosome protein changes but may not necessarily be associated with exosome quantity. Importantly, with the use of cell-type specific exosome reporter mice, we also showed that DREADD-mediated neuronal excitation significantly increases *in vivo* spreading of cortical neuronal exosomes. These *in vitro* and *in vivo* findings provide important insights about how glutamatergic synaptic transmission may stimulate somatodendritic secretion and spreading of exosomes in the CNS. As neuronal exosomes have been implicated in mediating the spreading of specific disease-associated protein aggregates in neurodegenerative diseases (Howitt and Hill, 2016), abnormal neuronal activity - especially hyperexcitability, commonly associated with these neurodegenerative diseases, including AD (Targa Dias Anastacio et al., 2022) and ALS (Xie et al., 2023) - may facilitate spreading of protein aggregates by promoting spreading of neuronal exosomes, thus contributing to disease pathology. In addition, our immunogold GFP labeling also frequently revealed axonal terminal localization and budding of hCD63-GFP^+^ vesicles in dopaminergic neurons, potentially suggesting axon-dependent release of exosomes, which is quite opposite to their preferential somatodendritic localization in forebrain excitatory neurons(Men et al., 2019). However, given the highly heterogenous neuronal subtypes and functions, such diverse subcellular localization of ILVs and release is not unexpected. Additional immunoEM analysis of ILVs/exosomes *in situ* from specialized neuronal subtypes and regions will provide further anatomical evidence of how exosomes operate in comparison to typical synaptic vesicles.

Neuronal exosomes in human CSF and plasma, either from control or neurological disease patients, have been commonly enriched and detected by immunoprecipitation of selective surface proteins (Hill, 2019; You et al., 2023), suggesting that neurons constantly secrete exosomes. By generating double reporter mice that concurrently and selectively label both specialized neuronal subtypes (Hb9^+^ MNs or DAT^+^ dopaminergic neurons) and exosomes, we observed extracellularly localized exosomes as early as in embryonic stages. Developmentally immature neurons have very primitive and less specialized morphology at embryonic and early postnatal stages. Neuronal axons still undergo pathfinding with mostly irregular spontaneous firing and the neural connectivity has not formed yet to have robust and directional synaptic transmission. Thus, neuronal secretion of exosomes may serve as important alternative signals to communicate with various CNS cell types during early CNS development. Consistently, previous *in vitro* studies have shown that human iPSC-derived neurons secrete exosomes to promote neurogenesis and circuit assembly (Sharma et al., 2019). Embryonic neurons also secrete EphB2^+^ exosomes to induce growth cone collapse by interacting with ephrinB1, which contributes to repulsive axon guidance (Gong et al., 2016). Developing neurons-derived exosomes control dendritic spine development through regulation of HDAC2 signaling (Zhang et al., 2021). In parallel, our small RNA-Seq datasets also showed abundant and diverse miRs in neuronal exosomes, some of which are predicted to preferentially bind to cell-type specific mRNAs in different CNS cell types. The roles of such selective intercellular miR-mediated genetic regulation in CNS development will be further investigated in future studies.

Although glutamate stimulation significantly promotes secretion of neuronal exosomes, our small RNA-Seq results showed that it has a very modest effect on neuronal exosomal profile of miRs. By further examining the cellular and exosomal abundance of selected top miRs, we showed that miRs are differentially enriched in exosomes. In particular, although miR-124-3p is 100 times less abundant in neurons than miR-9-5p, its abundance in exosomes is close to 50% of miR-9-5p in exosomes, resulting in > 20 times more enrichment than miR-9-5p. This is consistent with previous observations that various RNA-binding proteins, from hnRNPA2B1 and SYNCRIP to Ago2, may selectively bind certain miRs to carry them into exosomes for enrichment (McKenzie et al., 2016; Santangelo et al., 2016; Villarroya-Beltri et al., 2013). On the other hand, other miRs, such as miR-125b-5p, is highly abundant in both neurons and neuronal exosomes, suggesting that it may enter exosomes by non-selective RNA-binding mechanisms (Xia et al., 2022). As exosomal miRs may preferentially target on different CNS cell types based on predicted mRNA binding, a better understanding of how RNA-binding proteins mediate miR sorting into ILVs will help elucidate exosome miR-targeted neuron to other CNS cell-type communication.

## Acknowledgments

Imaging was performed with the assistance of the Tufts Center for Neuroscience Research. EM was performed with the assistance of the Harvard Medical School EM Core Facility. Small RNA library preparation and sequencing was carried out with the assistance of Tufts University Core Facility Genomics Core (supported by NIH shared instrumentation grant 1S10OD032203-01). This work was supported by NIH grants R01NS118747, R01NS125490, and R01AG078728 (YY).

## Author contributions

MB performed the majority of experiments in this study, analyzed data, and wrote part of the manuscript. QL performed DREADD injection, c-Fos immunostaining, and hCD63-GFP image analysis. FEM helped with c-Fos immunoreactivity analysis and performed VTA injection and immunoEM analysis. KER and KK performed miR qPCR, miR to mRNA target analysis, and quantification of hCD63-GFP^+^ immunogold signals; RJ generated triple positive mice and performed hCD63-GFP image analysis form triple positive mice; HW and JW performed bioinformatic analysis on small RNA sequencing results; YY designed overall study, analyzed data, and wrote the manuscript.

